# CAMISIM: Simulating metagenomes and microbial communities

**DOI:** 10.1101/300970

**Authors:** Adrian Fritz, Peter Hofmann, Stephan Majda, Eik Dahms, Johannes Dröge, Jessika Fiedler, Till R. Lesker, Peter Belmann, Matthew Z. Demaere, Aaron E. Darling, Alexander Sczyrba, Andreas Bremges, Alice C. Mchardy

**Affiliations:** Computational Biology of Infection Research, Helmholtz Centre for Infection Research, 38124 Braunschweig, Germany; Formerly Department of Algorithmic Bioinformatics, Heinrich-Heine University Düsseldorf, 40225 Düsseldorf, Germany; German Center for Infection Research (DZIF), partner site Hannover-Braunschweig, 38124 Braunschweig, Germany; Center for Biotechnology and Faculty of Technology, Bielefeld University, 33615 Bielefeld, Germany; The ithree institute, University of Technology Sydney, Sydney, New South Wales, Australia

## Abstract

Shotgun metagenome data sets of microbial communities are highly diverse, not only due to the natural variation of the underlying biological systems, but also due to differences in laboratory protocols, replicate numbers, and sequencing technologies. Accordingly, to effectively assess the performance of metagenomic analysis software, a wide range of benchmark data sets are required. Here, we describe the CAMISIM microbial community and metagenome simulator. The software can model different microbial abundance profiles, multi-sample time series and differential abundance studies, includes real and simulated strain-level diversity, and generates second and third generation sequencing data from taxonomic profiles or *de novo.* Gold standards are created for sequence assembly, genome binning, taxonomic binning, and taxonomic profiling. CAMSIM generated the benchmark data sets of the first CAMI challenge. For two simulated multi-sample data sets of the human and mouse gut microbiomes we observed high functional congruence to the real data. As further applications, we investigated the effect of varying evolutionary genome divergence, sequencing depth, and read error profiles on two popular metagenome assemblers, MEGAHIT and metaSPAdes, on several thousand small data sets generated with CAMISIM. CAMISIM can simulate a wide variety of microbial communities and metagenome data sets together with truth standards for method evaluation. All data sets and the software are freely available at: https://github.com/CAMI-challenge/CAMISIM

## INTRODUCTION

Extensive 16S rRNA gene amplicon and shotgun metagenome sequencing efforts have been and are being undertaken to catalogue the human microbiome in health and disease [1, 2] and to study microbial communities of medical, pharmaceutical, or biotechnological relevance [3–8]. We have since learned that naturally occurring microbial communities cover a wide range of organismal complexities – with populations ranging from half a dozen to likely tens of thousands of members – can include substantial strain level diversity, and vary widely in represented taxa [9–12]. Analyzing these diverse communities is challenging.

The problem is exacerbated by use of a wide range of experimental setups in data generation and the rapid evolution of short- and long-read sequencing technologies [13, 14]. Owing to the large diversity of generated data, the possibility to generate realistic benchmark data sets for particular experimental setups is essential for assessing computational metagenomics software.

CAMI, the initiative for the Critical Assessment of Metagenome Interpretation, is a community effort aiming to generate extensive, objective performance overviews of computational metagenomics software [15]. CAMI organizes bench-marking challenges and encourages the development of standards, and reproducibility in all aspects, such as data generation, software application, and result interpretation [16].

We here describe CAMISIM, which was originally written to generate the simulated metagenome data sets used in the first CAMI challenge. It has since been extended into a versatile and highly modular metagenome simulator. We demonstrate the usability and utility of CAMISIM with several applications. We generated complex, multi-replicate benchmark data sets from taxonomic profiles of human and mouse gut microbiomes [1, 17]. We also simulated thousands of small “minimally challenging metagenomes” to characterize the effect of varying sequencing coverage, evolutionary divergence of genomes, and sequencing error profiles on the popular MEGAHIT [18] and metaSPAdes [19] assemblers.

## THE CAMISIM SOFTWARE

CAMISIM allows customization of many properties of the generated communities and data sets, such as the overall number of genomes (community complexity), strain diversity, the community genome abundance distributions, sample sizes, the number of replicates, and sequencing technology used. For setting these options, a configuration file is needed, which is described in the Supplement. Simulation with CAMISIM has 3 stages (Figure 1):

1. design of the community, which includes selection of the community members and their genomes, and assigning them relative abundances,
2. metagenome sequencing data simulation, and
3. postprocessing, where the binning and assembly gold standards are produced.

**Fig. 1.**
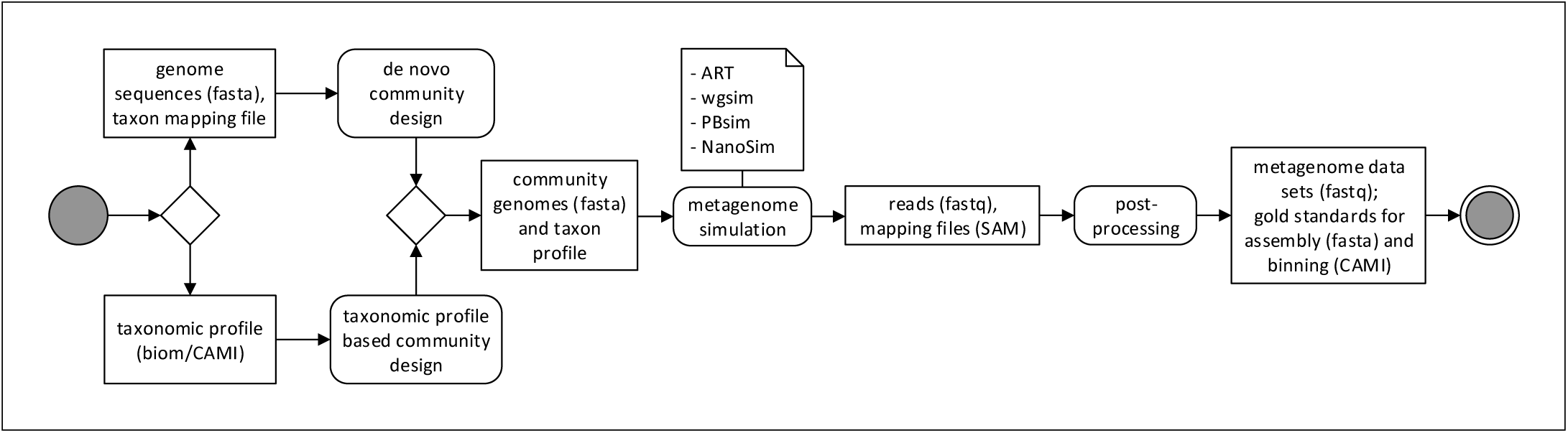
UML diagram of the CAMISIM workflow. CAMISIM starts with the “Community Design” step, which can either be *de novo*, requiring a taxon mapping file and reference genomes or based on a taxonomic profile. This step produces a community genome and taxon profile which is used for the metagenome simulation using one of currently four read simulators (ART, wgsim, PBsim, NanoSim). The resulting reads and bam-files mapping the reads to the original genomes, are used to create the gold standards before all the files can be anonymized and shuffled in the post-processing step.

### Community design

In this step, the community genome abundance profiles, called *P_out_*, are created. These also represent the gold standard for taxonomic profiling and, from the strain to the superkingdom rank, specify the relative abundances of individual strains (genomes) or their parental taxa in percent. In addition, a genome sequence collection for the strains in *P_out_* is generated. Both *P_out_* and the genome sequence collection are needed for the metagenome simulation in step 2. The taxonomic composition of the simulated microbial community is either determined by user-specified taxonomic profiles or generated *de novo* by sampling from available genome sequences.

#### Profile-based design

Taxonomic profiles can be provided in BIOM (Biological Observation Matrix) format [20]. With input profiles, the NCBI complete genomes [21] are used as the sequence collection for creating metagenome data sets. Optionally, the user can choose to also include genomes marked as “scaffold” or “contig” by the NCBI. Input genomes are split at positions with multiple occurrences of ambiguous bases, such that no reads spanning contig borders within larger scaffolds are simulated.

Profiles can include bacterial, archaeal and eukaryotic taxa, as well as viruses. The taxonomic identifiers of BIOM format are interpreted as free text scientific names and are mapped to NCBI taxon IDs (algorithm in the supplement). The so generated input profile *P_in_* specifies pairs (*t*, *ab_t_*) of taxon IDs *t* and taxon abundances *ab_t_* ∈ ℝ_≥0_. The profile taxa are usually defined at higher ranks than strain and thus have to be mapped approximately to the genome sequence collection for creating *P_out_.*

Given an ordered list of ranks *R* = (*species*, *genus*, *family*, *order*, *class*, *phylum*, *superkingdom*), CAMISIM requires as an additional parameter a highest rank *r_max_* ∈ *R*. We define the binary operator ≺, based on the ordering of the ranks in *R*. Given two ranks *r_i_*, *r_j_* ∈ *R*, we write *r_i_* ≺ *r_j_*, if *r_i_* appears before *r_j_* in *R* and we say *r_i_* is below *r_j_*. Related complete genomes are searched for all ranks below *r_max_*. By default this is the *family* rank. Another parameter is the maximum number of strains *m* that are included for an input taxon in a simulated sample.

To create *P_out_* from *P_in_*, the following steps are performed: Let *G_in_* be the set of taxon IDs of the genome collection at the lowest annotated taxonomic rank, usually *species* or *strain.* For all *t* ∈ *G_in_*, the reference taxonomy specifies a taxonomic lineage of taxon IDs (or undefined values) across the considered ranks in *R*. We use these to identify a collection of sets *F* = {*G_t_* | *t* = lineage taxon represented by ≥ 1 complete genome},

#### Algorithm 1. Creating a community genome abundance profile; *genome*-*select*(*F*, *P_in_*, *m*, *r_max_*)

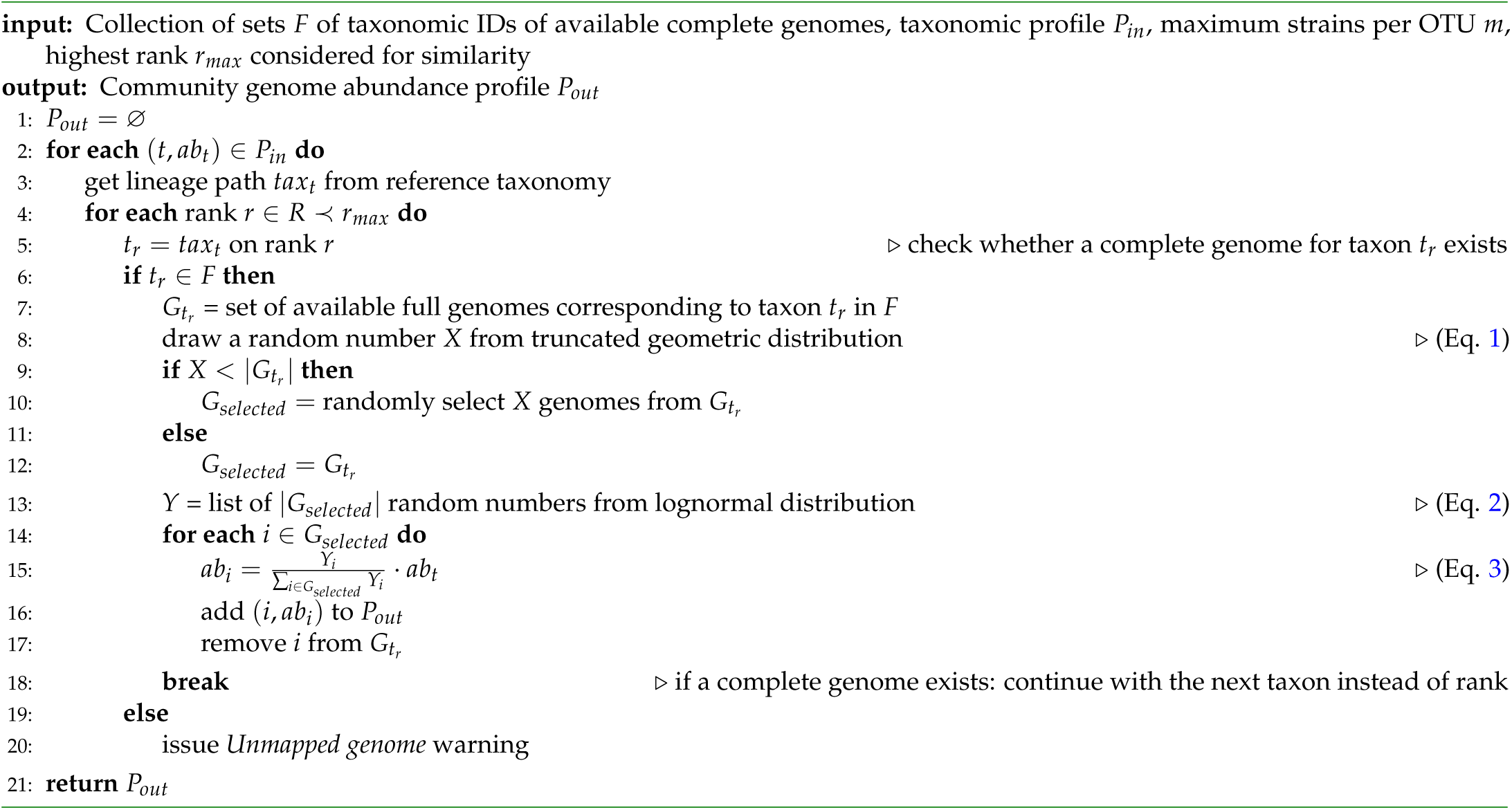

which specifies for each lineage taxon the taxon IDs of available genomes from the genome collection. *F* is used as input for Algorithm 1.

The algorithm retrieves for each *t* from the tuples (*t*, *ab_t_*) ∈ *P_in_* the lineage path *tax_t_* across the ranks of *R* (lines 2–3). Moving from the *species* to the highest considered rank, *r_max_*, the algorithm determines whether for a lineage taxon *t_r_* at the considered rank *r* a complete genome exists, that is, whether *G_t_* ≠ Ø for *t* = *t_r_* (lines 4–5). If this is the case, the search ends and *t_r_* is considered further (line 6). If no complete genome is found for a particular lineage, the lineage is not included in the simulated community, and a warning is issued (line 20). Next, the number of genomes *X* with their taxonomic IDs *t_r_* to be added to *P_out_* is drawn from a *truncated geometric distribution* (Eq. 1, line 8) with a mean of 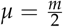 and the parameter *k* restricted to be less than *m*.

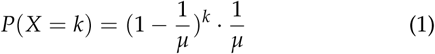

If | *G_t_r__* | is less than *X*, *G_t_r__* is used entirely as *G_selected_*, the genomes of *t_r_* that are to be included in the community. Otherwise *X* genomes are drawn randomly from *G_t_r__* to generate *G_selected_* (lines 9–12). It is optional to use genomes multiple times, by default the selected genomes *g* ∈ *G_selected_* are removed from *F*, such that no genome is selected twice (line 17). Based on the taxon abundances *ab_t_* from *P_in_*, the abundances *ab_i_* of the selected taxa *i* ∈ *G_selected_* for *t* are then inferred. First, random variables *ϒ_i_* are drawn from a configurable lognormal distribution, with by default mean *μ* = 1 and variance *σ* = 2 (Eq. 2) and then the *ab_i_* are set (Eq. 3; lines 13–15). Finally, the created pairs (*i*, *ab_i_*) are added to *P_out_* (line 16) and *P_out_* is returned (line 21).

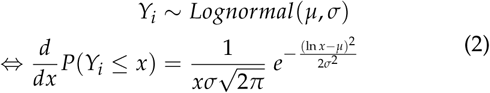

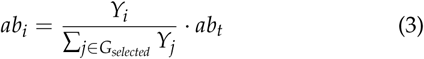

#### De novo design

A genome sequence collection to sample and a mapping file have to be specified. The mapping file defines for each genome a taxonomic ID (per default from the NCBI taxonomy), a novelty category and an Operational Taxonomic Unit (OTU) ID. Grouping genomes into OTUs is required for sampling related genomes, to increase strain-level diversity in the simulated microbial communities. The novelty category reflects how closely a query genome is related to draft or complete genomes in a genome sequence reference collection. This is used to maximize the spread of selected genomes across the range of taxonomic distances to the genome reference collection, such that there are genomes included of “novel” strains, species or genera. This distinction is relevant for evaluating reference-based taxonomic binners and profilers, which may perform differently across these different categories. The user can manually generate the mapping file as described in the supplement or in [15].

If controlled sampling of strains is not required, every genome can be assigned to a different OTU ID. If no reference based taxonomic binners or profilers are to be evaluated, or the provided genome sequence collection does not vary much in terms of taxonomic distance to publicly available genomes used as references for these programs, all genomes can be assigned the same novelty category.

In addition, the number of genomes *g_real_* to be drawn from the input genome selection and the total number of genomes *g_tot_* for the community genome abundance profile *P_out_* have to be specified. The *g_real_* real genomes are drawn from the provided genome sampling collection. An equal number of genomes is drawn for every novelty category. If the number of genomes for a category is insufficient, proportionately more are drawn from others. In addition, CAMISIM simulates *g_sim_* = *g_tot_* – *g_real_* genomes of closely related strains from the chosen real genomes in total. These genomes are created with an enhanced version of sgEvolver [22] (Supplementary Methods) from a subset of randomly selected real genomes. Given *m*, the maximum number of strains per OTU, up to *m* – 1 simulated strain genomes are added *per genome.* The exact number of genomes *X* to be simulated for a selected OTU is drawn from a geometric distribution with mean *μ* = 0.3^−1^ (Eq. 1) This procedure is repeated until *g_sim_* related genomes have been added to the community genome collection, comprising *g_tot_* = *g_real_* + *g_sim_* genomes [15].

Next, community genomes are assigned abundances. The relevant user-defined parameters for this step are the sample type and the number of samples *n*. In addition to single samples, multi-sample data sets (with differential abundances, replicates or time series) have become widely used in real sequencing studies [23–26], also due to their utility for genome recovery using covariance-based genome binners such as CONCOCT [27] or MetaBAT [28]. Several options for creating multi-sample metagenome data sets with these setups are provided:

1. If simulating a *single sample data set*, the relative abundances are drawn from a log-normal distribution, which is commonly used to model microbial communities [29–31]. By default, the mean is set to 1 and the standard deviation to 2 (Eq. 2). The two parameters of the lognormal distribution can be changed. Setting the standard deviation *σ*^2^ to 0 results in a uniform distribution.
2. The *differential abundance mode* models a community sampled multiple times after the environmental conditions or the DNA extraction protocols (and accordingly the community abundance profile) have been altered. This mode creates *n* different lognormally (Eq. 2) distributed genome abundance profiles.
3. Metagenome data sets with multiple samples with very similar genome abundance distributions can be created using the *replicates mode.* Having multiple replicates of the same metagenome has been reported to improve the quality for some metagenome analysis software, such as for genome binners [23, 27, 32, 33]. Based on an initial log-normal distribution *D*_0_, *n* samples are created by adding Gaussian noise to this initial distribution (Eq. 4). The Gaussian term accounts for all kinds of effects on the genome abundances of the metagenomic replicates including, but not limited to, different experimenters, different place of extraction, or other batch effects.

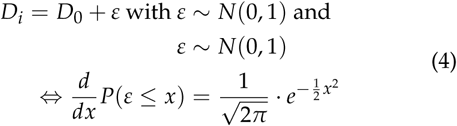
4. *Time series* metagenome data sets with multiple related samples can be created. For these, a Markov model-like simulation is performed, with the distribution of each of the *n* samples (Eq. 5) depending on the distribution of the previous sample plus an additional either lognormal (Eq. 2) or Gaussian (Eq. 4) term. This emulates the natural process of fluctuating abundances over time and ensures that the abundance changes to the previously sampled metagenome do not grow very large.

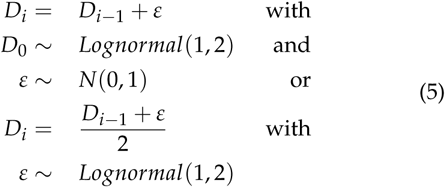

### Metagenome simulation

Metagenome data sets are generated from the genome abundance profiles of the community design step. For each genome-specific taxon *t* and its abundance (*t*, *ab_t_*) ∈ *P_out_*, its genome size *s_t_*, together with the total number of reads *n* in the sample, determines the number of generated reads *n_t_* (Eq. 6). The total number of reads *n* is the overall sequence sample size divided by the mean read-length of the utilized sequencing technology.

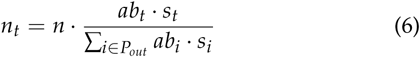

By default, ART [34] is used to create Illumina 2 × 150 bp paired-end reads with a HiSeq 2500 error profile. The profile has been trained on MBARC-26 [35], a defined mock community that has already been used to benchmark bioinformatics software and a full-length 16S rRNA gene amplicon sequencing protocol [36, 37], and is distributed with CAMISIM. Other ART profiles, such as the one used for the first CAMI challenge, can also be used. Further available read simulators are wgsim (https://github.com/lh3/wgsim, originally part of SAMtools [38]) for simulating error-free short reads, pbsim [39] for simulating Pacific Biosciences data and nanosim [40] for simulating Oxford Nanopore Technologies reads. The read lengths and insert sizes can be varied for some simulators.

For every sample of a data set, CAMISIM generates FASTQ files and a BAM file [38]. The BAM file specifies the alignment of the simulated reads to the reference genomes.

### Gold standard creation and postprocessing

From the simulated metagenome data sets – the FASTQ and BAM files – CAMISIM creates the assembly and binning gold standards. The software extracts the perfect assembly for each individual sample, and a perfect co-assembly of all samples together by identifying all genomic regions with a coverage of at least one using SAMtools’ mpileup and extracting these as error-free contigs. This gold standard does not include all genome sequences available for the simulation, but the best possible assembly of their sampled reads.

CAMISIM generates the genome and taxon binning gold standards for the reads and assembled contigs, respectively. These specify the genome and taxonomic lineage that the individual sequences belong to. All sequences can be anonymized and shuffled (but tracked throughout the process), to enable their use in benchmarking challenges. Lastly, files are compressed with gzip and written to the specified output location.

## RESULTS

### Comparison to the state-of-the-art

We tested seven simulators and compared them to CAMISIM (Table 1). All generate Illumina data and some – NeSSM [44], BEAR [45], FASTQSim [46] and Grinder [47] – also use a taxonomic profile. Novel and unique to CAMISIM is the ability to simulate long-read data from Oxford Nanopore, of hybrid data sets with multiple sequencing technologies and multi-sample data sets, such as with replicates, time series or differential abundances. Grinder [47] can also create multiple samples, but only with differential abundances. In addition, CAMISIM creates gold standards for assembly (single sample assemblies and multi-sample co-assemblies), for taxonomic and genome binning of reads or contigs and for taxonomic profiling. Finally, CAMISIM can evolve multiple strains for selected input genomes, and allows specification of the degree of real and simulated intra-species heterogeneity within a data set.

**Table 1.**
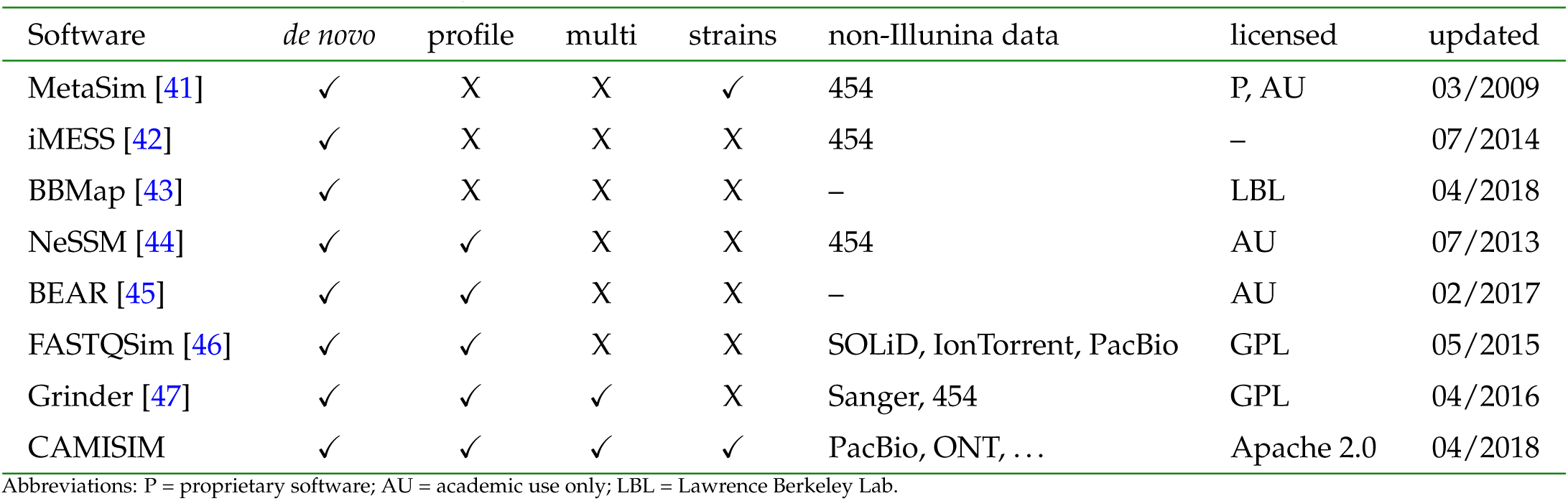
Properties of popular metagenome sequence simulators. The table shows if an abundance profile can be generated by the simulator *de novo* and if an existing *profile* of a microbial community can be used as input. Further inspected features are the ability to simulate *multi*-sample data sets, *strains*, and *non*-*Illumina data* (e.g. long reads). Lastly, the table states if and how a software is *licensed*, and the date it was last recently *updated*.

### Effect of data properties on assemblies

We created several thousand “minimally challenging” metagenome samples by varying one data property relevant for assembly, while keeping all others the same. Using these, we studied the effect of evolutionary divergence between genomes, different error profiles and coverage on the popular metaSPAdes [19], version 3.7.0, and MEGAHIT [18], versions 1.1.2 and 1.0.3, assemblers, to systematically investigate reported performance declines for assemblers in the presence of strain-level diversity, uneven coverage distributions and abnormal error profiles [15, 48, 49]. Both MEGAHIT and metaSPAdes work on de Bruijn graphs, which are created by splitting the input reads into smaller parts, the *k*-mers, and connecting two *k*-mers if they overlap by exactly *k*-1 letters. For every sequencing error *k* erroneous *k*-mers are introduced into the de Bruijn graph, which might substantially impact assembly (Figure 2).

**Fig. 2.**
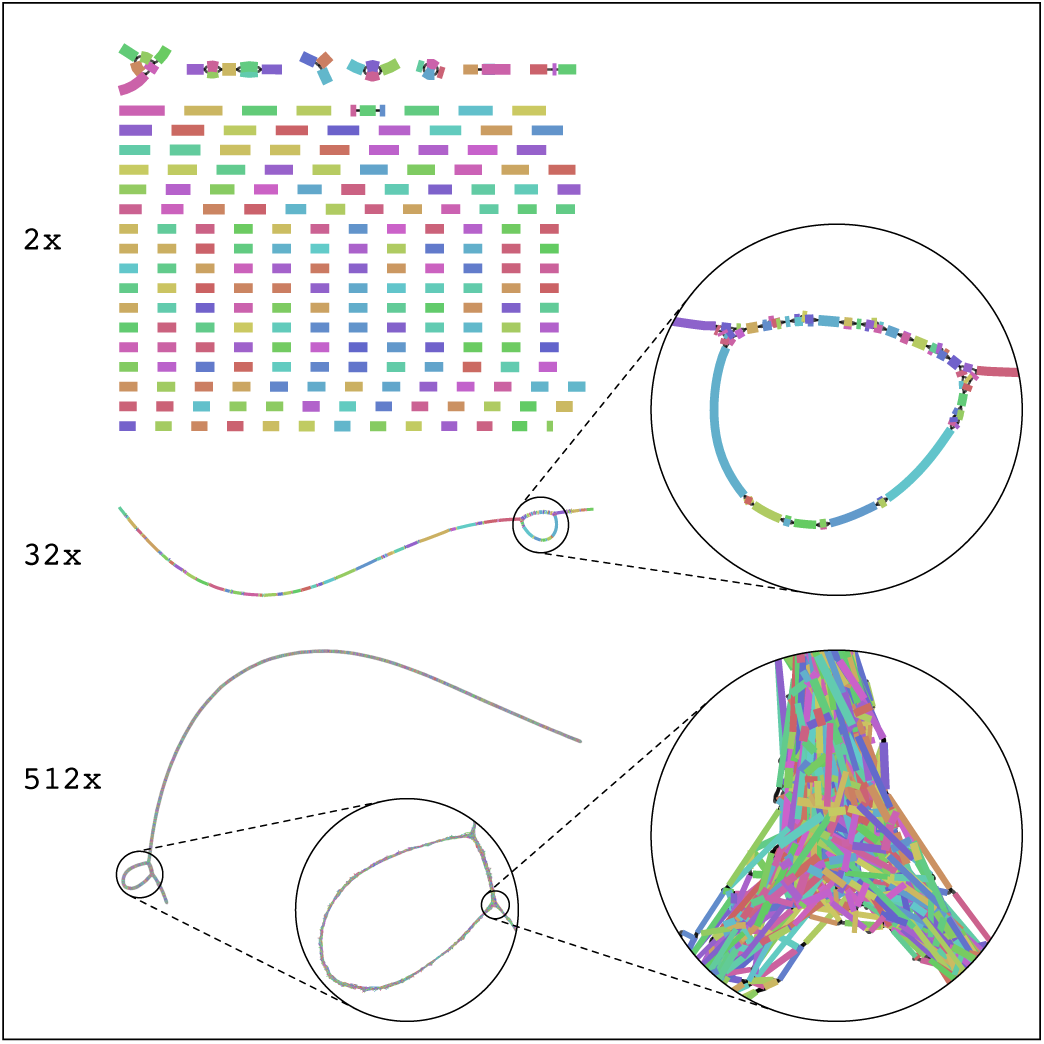
Assembly graphs become more complex as coverage increases. MEGAHIT assembly graphs (*k*=41) of an *E. coli* K12 genome for 2x, 32x, and 512x per-base coverage, respectively, visualized with Bandage [50]. For 2x coverage, the graph is disconnected and thus the assembly fragmented. With increasing coverage more and more unitigs can be joined, first resulting in a decent assembly for 32x coverage, but – due to sequencing errors adding erroneous edges to the graph – a fragmented assembly again for 512x coverage.

#### Varying genome coverage and sequencing error rates

We initially simulated samples from *Escherichia coli* K12 MG1655 with varying coverage and different error rates. Reads were generated at 512x genome coverage and subsampled stepwise by 50% until 2x coverage was reached, resulting in a sample series with 512, 256, 128, 64, 32, 16, 8, 4 and 2-fold coverage, respectively. Subsampling was employed to control variation in the sampling of different genomic regions. To assess the effect of sequencing errors, 4 read data sets were simulated; three using wgsim with uniform error rates of 0%, 2% and 5%, and one using ART with the CAMI challenge error profile (ART CAMI).

Both assemblers were run on these data sets with default options, except for the phred-offset parameter for metaSPAdes, which was set to 33. Both assemblers performed similar across all error rates and coverages, with assembly quality varying substantially with coverage (Figure 3). Performance on the data generated with the 5% error profile was worst throughout. This is an unrealistically high error profile for Illumina data [49] that software need not necessarily be adapted to handle well.

**Fig. 3.**
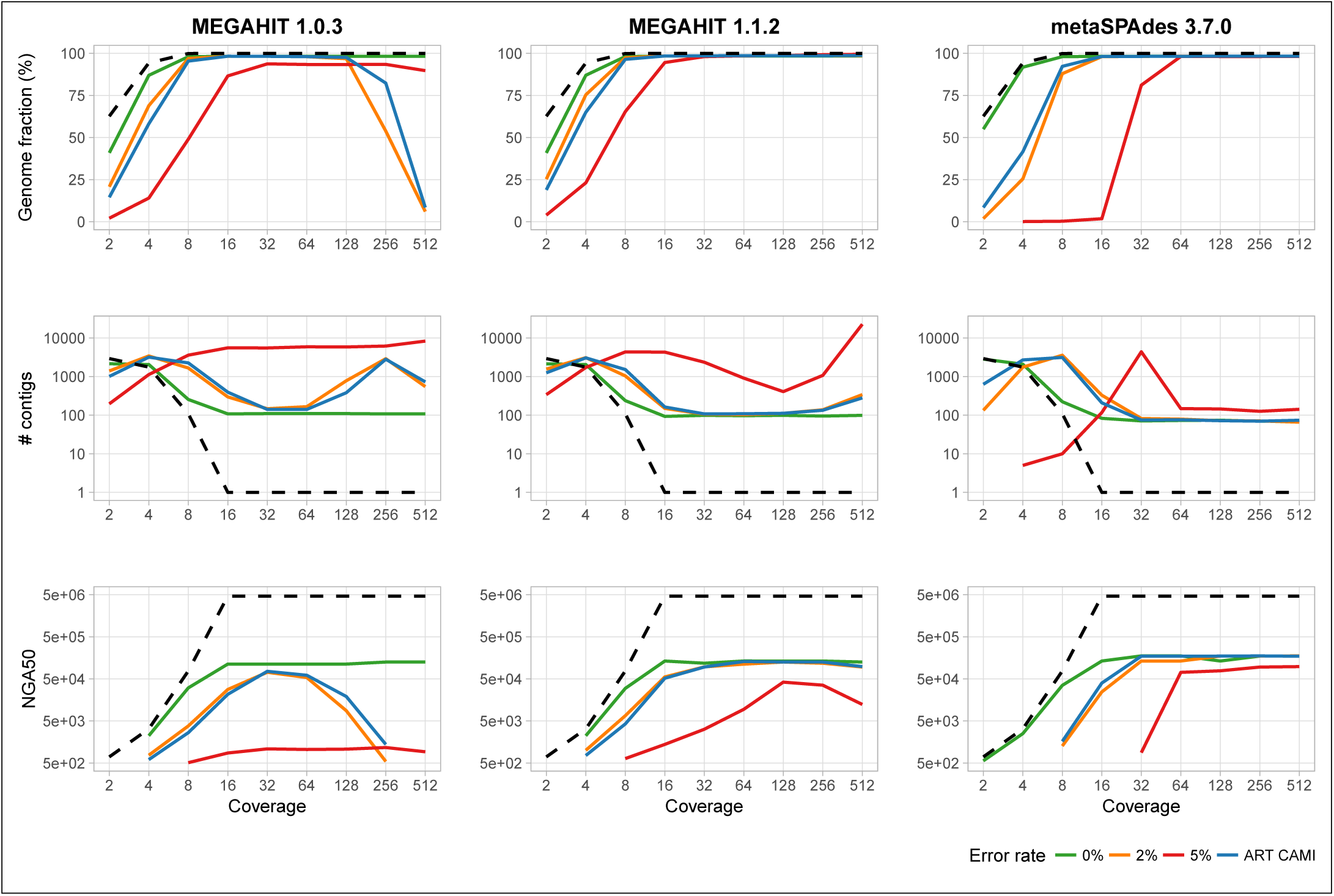
Coverage dependent assembly performance for MEGAHIT and metaSPAdes. Shown are the metrics, from top to bottom: Genome fraction in %, number of contigs, and NGA50 (as reported by QUAST [51]), for 0%, 2%, and 5% uniform error rate, and with the ART CAMI error profile compared to the best possible metrics (gold standard) on the ART CAMI profile (dashed black).

If coverage was low, assembly failed, generating a large number of small (low NGA50) contigs covering only a small genome portion (genome fraction) across all data sets, because of uncovered regions in the genomes. Sequencing errors (denoted *ε*) do not play a major role (Figure 2). The expected per-base error-rate *E_p_* = cov · *ε* (disregarding the biased errors in the short-read sequencing technologies) is far below 1 (*E_p_* ≪ 1). With increasing coverage, assembly improved consistently across the 0%, 2% and CAMISIM ART error profile data sets and both assemblers for all metrics (Figure 3), and reaching an early plateau by 8–16x coverage.

Notably, the performance of an earlier version of MEGAHIT (1.0.3) decreased substantially (declining genome fraction and NGA50) for more than 128x coverage, except for error-free reads. For instance, at 5% error rate, MEGAHIT, version 1.0.3, generated an exponential number of contigs at high coverages, which keeps the genome fraction artificially high. For these high coverages and error rates, we expect multiple errors at every position of the genome (*E_p_* ≫ 10). This creates de Bruijn graphs with many junctions and bubbles (Figure 2) which cannot easily be resolved and may lead to breaking the assembly apart and covering the same part of the genome with multiple, short, erroneous contigs (Figure 3).

#### Effect of evolutionary relatedness on assembler performances

We systematically investigated the effect of related strains on assembler performances across a wide range of taxa and evolutionary divergences, using the genomes of 152 species from the interactive tree of life iTol [52], which includes bacteria, archaea and eukaryotes. For each genome we evolved 19 related genomes without larger insertions and deletions and an Average Nucleotide Identity (ANI) between 90% and 99.5% to the original one using steps of 0.5%. For each of the 152 · 20 = 3,040 pairs of original and evolved genome sequences, we simulated single sample minimal metagenomes at equal genome abundances, with error-free reads at 50*x* coverage using wgsim. This constitutes good coverage for the analyzed assemblers, as shown in the previous section. For the resulting samples, variation in assembler performance should thus primarily be caused by differences in ANI.

The presence of closely related genomes substantially affected assembly quality (Figure 4). For up to 95% ANI, the assemblers restored high quality assemblies for both genomes. Between 95% and 99% ANI, the genome fraction and assembly size dropped substantially and contig numbers increased. This was the case if we allowed contigs to either map uniquely to one reference genome or to both, in case of multiple optimal mappings. For more closely related genomes, the number of contigs increased drastically and the assembly size continued to drop. The genome fraction remained high when considering non-unique mappings, but decreased for unique mappings: the explanation for this observed behavior is that for an ANI of more than 99%, assemblers produced consensus contigs of the two strains that mostly aligned similarly well to both reference genomes. This was the case for all 152 genomes and their evolved counterparts.

**Fig. 4.**
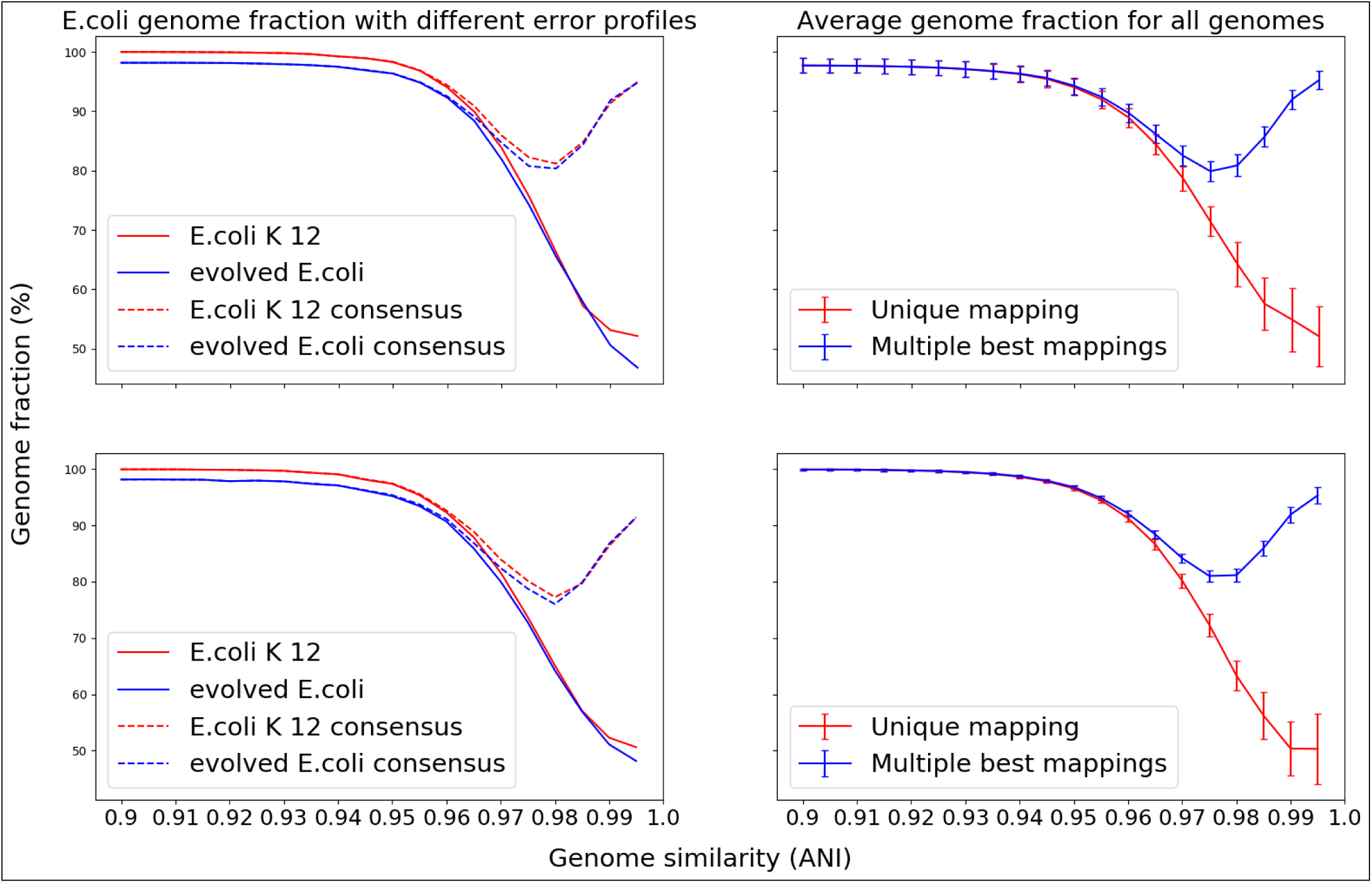
Genome fraction calculated using unique or multiple best mappings in case of ties to the community genome collection. Left: Genome fraction for the *E. coli* assembly created by MEGAHIT from error-free reads (top) and with ART CAMI error profile (bottom). Right: Average genome fraction and standard deviation for all original 152 iTol genomes created by MEGAHIT from error-free reads (top) and with ART CAMI error profile (bottom). Error bars denote 1x standard deviation.

### Simulating environment-specific data sets

To test the ability to create metagenome data of the human microbiome, we simulated metagenomes from taxonomic profiles of the Human Microbiome Project [9] for different body sites with CAMISIM. We selected 49 samples from the airways, gastrointestinal tract, oral cavity, skin and urogenital tract, with whole genome shotgun (WGS) and 16S rRNA gene amplicon sequence data available. We used the published qiime OTU table (https://hmpdacc.org/hmp/HMQCP/) to generate 5 Gb of simulated reads per sample with CAMISIM, resulting in a data set of 245 Gb of Illumina data, and of PacBio data, respectively. Only genomes tagged as “complete genomes” in the NCBI were considered in the data set generation. To decrease the chance of OTUs not being represented by a genome, the option of allowing multiple OTUs being represented by a single reference genome was turned on. This can be relevant for instance when due to sequencing errors in 16S rRNA data, individual community genomes are represented by multiple OTUs.

For a functional comparison of the simulated data with the original metagenome shotgun data, we inferred KEGG Ortholog family abundance profiles from the raw read data sets [53]. To this end, all reads were searched with Diamond v0.9.10 using its blastx command with default options [54] against the KEGG GENES database (release 77, species_prokaryotes, best-hit approach) and linked to KEGG Orthology (KO) via the KEGG mapping files. KO profile similarity between the simulated and original metagenome samples was calculated with Pearson’s correlation coefficient (PCC) and Spearman rank correlation (SRC), and visualized with non-metric multidimensional scaling (NMDS) [55].

For comparison we also created functional profiles with PICRUSt [56], using a prediction model generated from 3772 KEGG genomes and corresponding 16S rRNA gene sequences according to the PICRUSt “Genome Prediction Tutorial” (Supplementary Information). The PCC of the CAMISIM and original samples approached a striking 0.97 for body sites with high bacterial abundances and many sequenced genomes available, such as the GI tract and oral cavity, and still ranged from 0.72 to 0.91 for airways, skin and urogenital tract samples (Figure 5B). All PCCs were 7-30% higher than the PCC of PICRUSt with the original metagenome samples. Thus CAMISIM created metagenome samples functionally even closer to the original metagenome samples than the functional profiles created by PICRUSt. The higher PCC may also partly be due to the fact that the original and CAMISIM data were annotated by “blasting” reads versus KEGG, while the PICRUSt profiles were directly generated from KEGG genome annotations. The Spearman correlation of the simulated CAMISIM samples to the original metagenome samples was slightly lower than the PCC across all body sites, and very similar for CAMISIM and PICRUSt (0–6% improvement of CAMISIM over PICRUSt). These results demonstrate the quality of the CAMISIM samples.

**Fig. 5.**
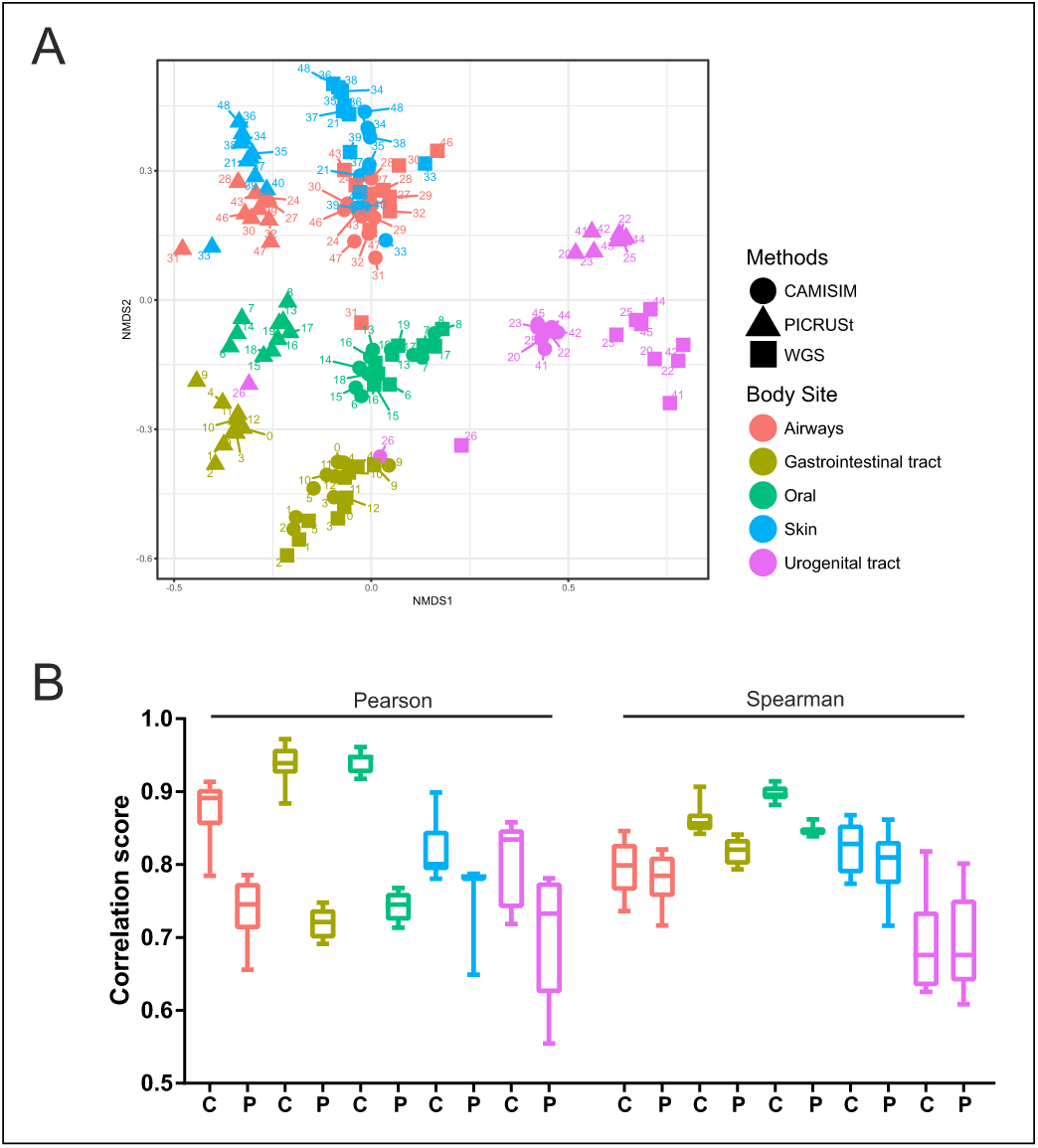
Comparison of CAMISIM and PICRUSt functional profiles for different body sites. (A) NMDS ordination of the functional predictions of individual samples by the different methods. The different body sites are color-coded and labeled with their sample number. The original WGS is denoted by squares, the CAMISIM result as circles and the PICRUSt result as triangles. (B) Mean and standard deviation of Pearson and Spearman correlation to original WGS samples per body site. C: CAMISIM; P: PICRUSt.

The NMDS plot (Figure 5A) showed a very distinct clustering of the CAMISIM and original WGS samples by body site, more closely than the original samples clustered with the PICRUSt profiles. Even though the urogenital tract samples did not cluster perfectly, the CAMISIM samples still formed a very distinct cluster close to the original one. Even outliers in the original samples were, at least partly, detected and correctly simulated (both original and simulated sample 26 of urogenital tract cluster most closely with the gastrointestinal tract microbiomes).

We also provide a multi-sample mouse gut data set for software developers to benchmark against. For 64 16S rRNA samples from the mouse gut [17], we simulated 5 Gb of Illumina and PacBio reads each. The mice were obtained from 12 different vendors and the samples characterized by 16S V4 amplicon sequencing (OTU mapping file in the supplement). Since for mouse gut only a few complete reference genomes were available, the “scaffold” quality for downloading genomes was chosen.

## DISCUSSION AND CONCLUSIONS

CAMISIM is a flexible program for simulating a large variety of microbial communities and metagenome samples. To our knowledge it possesses the most complete feature set for simulating realistic microbial communities and metagenome data sets. This feature set includes: simulation from taxonomic profiles as templates, inclusion of both natural and simulated strain level diversity, and modelling multi-sample data sets with different underlying community abundance distributions. Read simulators are included for short read (Illumina) and long read (PacBio, ONT) sequencing technologies, allowing the generation of hybrid data sets. This turns CAMISIM into a versatile metagenome simulation pipeline, as modules for new (or updated) sequencing technologies and emerging experimental setups can easily be incorporated.

We systematically explored the effect of specific data properties on assembler performances on several thousand minimally challenging metagenomes. While low coverage reduced assembly quality for both assemblers, MetaSPAdes and MEGAHIT performed generally well for medium to high coverages and different error profiles. Notably, MEGAHIT is computationally very efficient and overall performed well. As noted before [15, 57], assemblers had problems with resolving closely related genomes in our experiments. For an in-depth investigation, we systematically analyzed the effect of related strains on MEGAHIT’s performance across a wide range of taxa and evolutionary divergences. The average nucleotide identity (ANI) between two genomes is a robust measure of genome relatedness; an ANI value of 95% roughly corresponds to a 70% DNA-DNA reassociation value – a historical definition of bacterial species [58, 59]. For an pairwise ANI below 95%, the mixture of strains could be separated quite well and assembled into different contigs. For an ANI of more than 99%, consensus contigs of strains were produced that mostly aligned similarly well to either reference genome. In the “twilight zone” of 95–99% nucleotide identity, assembly performance dropped substantially and MEGAHIT’s inability to reliable phase strain variation resulted in many small (and often redundant) contigs. For IDBA-UD [60], another *de Bruijn* graph-based metagenome assembler, a similar pattern has been observed [61], indicating that such behavior is common to many assemblers.

Resolving strains from metagenome shotgun data is an open research question, though recently promising computational approaches were proposed [11, 62]. The hybrid long and short read simulated data sets we are providing for mouse gut and human body sites could enable the development of new appraoches for this task CAMISIM will facilitate the generation of further realistic benchmarking data sets to assess their performances. It can also be used to study the effect of experimental design (e.g. number of replicates, sequencing depth, insert sizes) or intrinsic community properties, such as taxonomic composition, community abundance distributions, and organismal complexities, on program performance. Due to the enormous diversity of naturally occurring microbial communities, experimental and sequencing technology setups used in the field, such explorations are required to determine the most effective combinations for specific research questions.

## SOFTWARE AND DATA AVAILABILITY

CAMISIM is implemented in Python 2.7 and available under the Apache 2.0 license. The software, config files, input genomes, and metadata are available at: https://github.com/CAMI-challenge/ CAMISIM and https://github.com/CAMI-challenge/CAMISIM-DATA.

The large human and mouse gut microbiome data sets (alongside the BIOM and config files from which they were created) are available at: https://data.cami-challenge.org/participate.

## AUTHORS’ CONTRIBUTIONS

AF, PH, SM, ED, JD, JF, MZD, AED, and AB implemented CAMISIM; AF and PB tested the software; AF, TRL, and AB performed the experiments; AF, TRL, AS, AB, and ACM interpreted the results; AF, PH, TRL, PB, AB, and ACM wrote the manuscript; AB and ACM conceived the experiments; ACM conceived and supervised the project. All authors read and approved the final manuscript.

## ACKNOWLEDGEMENTS

The authors thank the Isaac Newton Institute for Mathematical Sciences for its hospitality during the programme MTG, which was supported by EPSRC Grant Number EP/K032208/1, and Victoria Sack for generating Figure 1. This research project has been supported by the President’s Initiative and Networking Funds of the Helmholtz Association of German Research Centres (HGF) under contract number VH-GS-202.

